# Visual input dynamically changes responses to spatiotemporal tactile input patterns in S1 neurons

**DOI:** 10.1101/2022.11.30.518507

**Authors:** Sofie Skårup Kristensen, Henrik Jörntell

**Affiliations:** Neural Basis of Sensorimotor Control, Department of Experimental Medical Science, Lund University, Lund, Sweden

## Abstract

To understand how sensory events are represented in and perceived by the brain, one must understand how varying internal brain states affect neuronal decoding of sensory input. Recent studies indicate global state changes in the brain impact the representation of haptic events in neurons of the primary somatosensory cortex (S1). It could be argued that the manipulations used so far to alter the cortical circuitry behavior were artificial and not reflective of normal information processing in the neocortex. In the present study we therefore wanted to explore if natural visual stimulation also could impact the interpretation of given tactile input patterns. We recorded the unitary extracellular responses to a set of spatiotemporal tactile input patterns presented either alone or together with simultaneously multicolor flashing lights from a large number of neurons in parallel in the rat primary somatosensory cortex (S1). We found that the visual input, mildly but consistently altered the temporal spike outputs to tactile input patterns in S1 neurons. We argue that the visual input change the global cortical state to an extent that it affects the cortical representation of haptic events even within the S1 and that this is an indication that the cortical network in its information processing may be far more reliant on globally distributed network dynamics than traditionally thought.

## Introduction

When you grasp an object, you do not experience the sight of the object and the haptic sensation as separate events. Your brain integrates information from spatiotemporal activation patterns of photo -and mechanoreceptors into a single percept and your experience becomes holistic and continuous. Using feed-forward network models as an interpretational framework, cortical multisensory integration was originally thought to occur only in secondary heteromodal areas, after unimodal processing in primary cortical regions (Alsius et al., 2005; Busse et al., 2005; Koelewijn et al., 2010; Talsma & Woldorff, 2005). A number of studies has, however, shown that neurons in primary sensory regions respond to sensory input from other modalities (Alsius et al., 2005; Busse et al., 2005; Enander et al., 2019; Koelewijn et al., 2010; Talsma & Woldorff, 2005; Wallace et al., 2004; Zhou et al., 2020). Following the finding that neurons in primary sensory regions do not only have modality specific neurons, it was even suggested that all of neocortex may be essentially multisensory in the organization of its processing (Ghazanfar & Schroeder, 2006).

Other recent studies also indicate that the neocortex may be more densely interconnected globally than what has previously been assumed. For example, it has been shown that perturbation of the activity in remote cortical areas, i.e. focal stroke-like lesions (Wahlbom et al., 2019) or local circuitry perturbation through intracortical electrical microstimulation (Etemadi et al., 2022), both have profound effects on the information processing of tactile input patterns in neurons of the primary somatosensory cortex. Etemadi et al. (2022) showed that even when perturbation does not result in a measurable response in an individual neuron, it still results in dynamic modulations of the cortical circuitry globally and thereby impact the responses of the neuron. Not only does this further challenge the notion of “unimodal” and “multimodal” cortical regions, it also suggests that even if there is no measurable response in an S1 neurons to e.g. visual stimulation, its response to tactile stimulation might still be modified by a change in the global neocortical state. Such global state would in theory be modifiable by any activity change in any part of the neocortex, and for example simultaneous visual input would in this scenario be expected to modify the global interpretation of a tactile input pattern. Feed-forward network theories would suggest that this would primarily occur in higher order multimodal cortical areas, but here we wanted to investigate if such changes would be detectable already at the level of the unimodal area, i.e. the primary somatosensory cortex (S1).

In the present study we aim to explore whether natural visual stimulation can impact the interpretation of given tactile input patterns in the individual neurons of S1. We hypothesize that visual stimulation will cause global dynamic network changes and impact the patterns of neuronal spike responses to tactile stimulation in somatosensory cortex (S1). We explored this by providing multicolor flashing lights as visual inputs, while recording the extracellular spiking responses of S1 neurons to complex spatiotemporal tactile input patterns, reminiscent of tactile input patterns that could arise in haptic interactions. As tactile input, we used an electrotactile interface by which we could deliver different specific spatiotemporal activation patterns. This method generates exactly reproducible patterns of tactile afferent activation, which poses an advantage compared to natural mechanical activation of the skin in that it reduces the variability in skin sensor activation, thus improving the precision of the analysis of the cortical responses. The spatiotemporal patterns were originally developed to deliver sensor activation imitating active touch and are suitable for quantification and statistical analysis of neural decoding (Oddo et al., 2017). As visual input we used flashing LEDs because these have been shown to induce strong responses in V1 (Griffen et al., 2017).

## Materials and methods

### Surgical procedure

Adult Sprague-Dawley rats (N=4, male sex, weight 306–420 g) were prepared and maintained under anesthesia in the same way as (Etemadi et al., 2022; Norrlid et al., n.d.). General anesthesia was induced with a mixture of ketamine/xylazine (ketamine: 40 mg/kg and xylazine: 4 mg/kg). The mixture was injected intraperitoneally while the rat was sedated with a mixture of air and isoflurane gas (3%) for 1-2 minutes. To maintain anesthesia, Ringer acetate and glucose mixed with anesthetic (ketamine and xylazine in a 20:1 ratio, delivered at a rate of ~5 mg/kg/h ketamine) was continuously infused through an intravenous catheter in the right femoral vein. The catheter was inserted by making an incision in the inguinal area of the hindlimb. We removed a small part of the skull (~2 x mm) to expose the somatosensory (S1) cortex. After inserting the Neuropixel probe, the exposed brain area was covered in a thin layer of agarose (1%) to prevent dehydration of the brain. To ensure a light level of anesthesia that would still prevent the animal from feeling pain, the hind paws were regularly pinched in different areas to verify the absence of withdrawal reflexes. Furthermore, the presence of desynchronized brain activity was continuously monitored. The anesthetic mixture was chosen because it has been reported to preserve the sequential order of neuronal recruitment at short time spans in stimulation evoked responses (Luczak & Barthó, 2012). Furthermore, light anesthesia minimizes brain activity noise caused by thought processes and uncontrollable movements, which would otherwise be a confounding factor for this analysis and would have given less consistent results. Animals were sacrificed after the end of the experiment with an overdose of pentobarbital.

### Stimulation

Because mechanoreceptors are located in movable tissue, repeated interactions between the same object and the same part of the skin, will give rise to different spatiotemporal activation patterns of the sensors. To overcome this variability, haptic stimulation was delivered as spatiotemporal patterns of electrical skin activation through four pairs of intracutaneous needle electrodes (channel 1-4). The needle electrodes were inserted into the volar side of the second digit of the forepaw with 2-3 mm distance between each pair. The stimulation patterns were delivered as pulses with intensities of 0.5 mA and durations of 0.14 ms (DS3 Isolated Stimulator, Digitimer, UK). Four different types of tactile input patterns were delivered in pre-defined random order, either alone or with simultaneous visual stimulation. Pattern durations lasted between 200 ms and 340 ms and were separated by about 1.8 s (random intervals). The time interval of 1.8 s was chosen because Oddo et al., (2017) showed that neurons continue to respond to stimulation up to 700 ms after the offset of a stimulation and that an interval of 1.8 s allows the cortical activity to settle in between stimulations. Each pattern was repeated 70 times during a protocol (35 times without visual stimulation, 35 times with visual stimulation), and protocols were repeated 6-10 times, hence providing up to 350 repetitions of each stimulus combination for each neuron recorded. The patterns mimic expected tactile afferent activation patterns for four different types of mechanical skin-object interactions and were developed by Oddo et al., (2017) using an artificial fingertip equipped with four neuromorphic sensors. The sensorized fingertip was moved against four objects with different roundness (illustrated in **Figure 1D**) and generated spiking output with a spiking neuron model mimicking a slow- and a fast-adapting mechanoreceptor, respectively. This resulted in eight different spatiotemporal activation patterns of the four sensors (slow and fast for each sensor) of which only the four patterns resembling the fast-adapting mechanoreceptor are used in the present paper. Each electrode pair in the rat skin corresponds to a spike output from one of the sensors generated by the object-sensor interface. The four spatiotemporal activation patterns are labelled according to the curvature of the object (“infinity” means flat) and the adaptation rate of the simulated skin receptors (F, fast; or S, slow).

**Figure 1.**
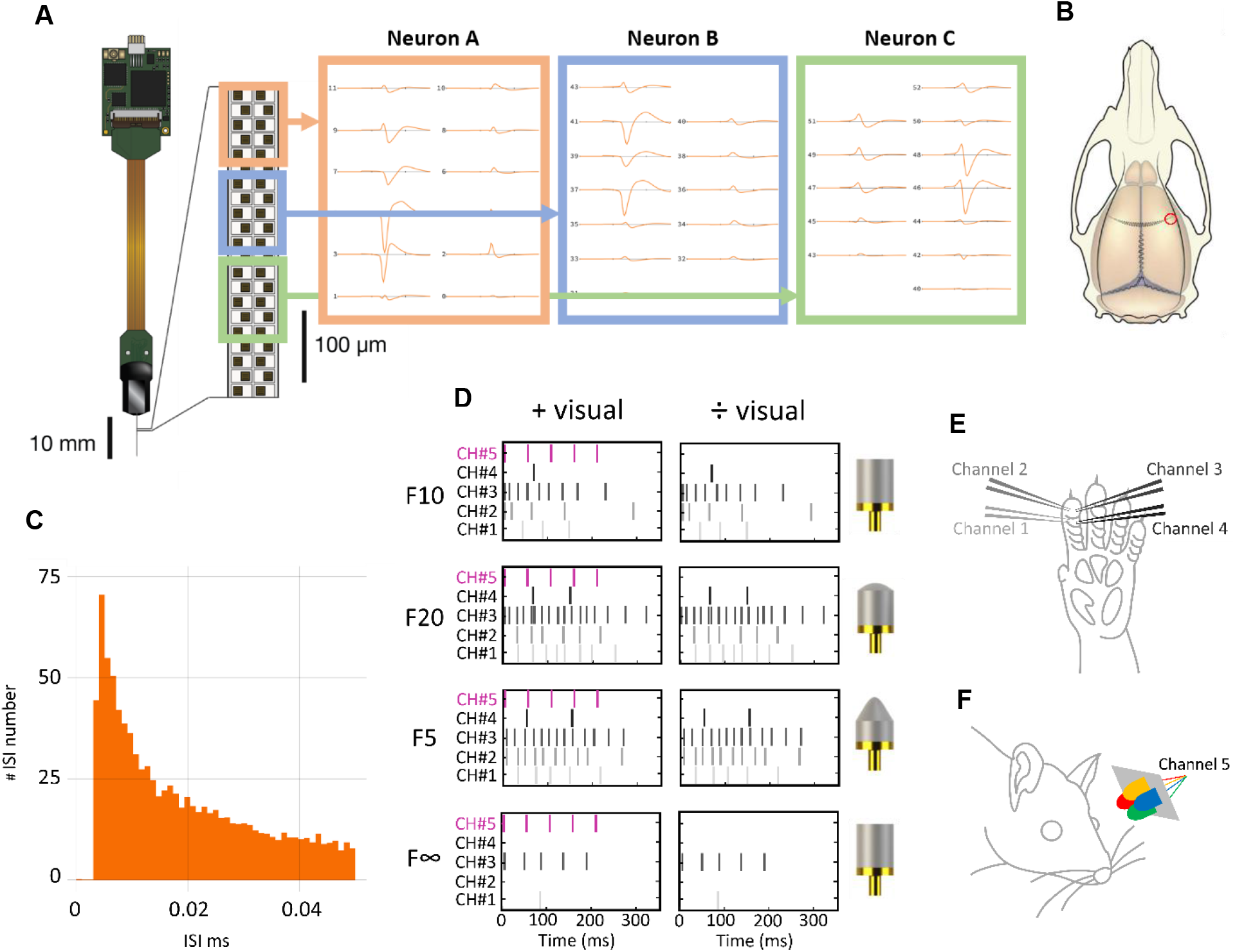
Recording and stimulations. **(A)** Illustration of the Neuropixel probe on the left. To the right are averaged spikes identified by the Matlab library Kilosort2.5 as three separate units. **(B)** Illustration of recording site S1 marked in red. **(C)** Screen print from the open-source Python library Phy of the inter-spike-interval (ISI) histogram for one sample neuron. **(D)** Illustration of spatiotemporal activation patterns (channel 1-4) with and without flashing lights from the LEDs (channel 5). To the right is an illustration of the four different surfaces the tactile patterns resemble the touching of. **(E)** Illustration of the four pairs of needle electrodes in the left forepaw of the rat. **(F)** Illustration of custom-made apparatus with four ascending LEDs in red, blue, green, and yellow.

Because we wanted to investigate the effect of visual stimulation on the cortical representation of tactile stimulation patterns, we did not have the same requirements for the visual stimulation to resemble anything naturalistic or meaningful. We wanted simple, reproducible visual stimulation strong enough to induce responses besides visual parts of cortex. Therefore, visual stimulation was delivered as pulses passed through a custom-made apparatus consisting of four ascending 7000 mcd LEDs. This type of visual stimulation has previously been shown to induce strong responses in V1 neurons (Griffen et al., 2017). The apparatus was positioned at a 30 ° viewing angle, 4-5 cm from the left eye and could deliver 1mW of power. 5 flashes were delivered with 5 ms onset and 50 ms offset, always starting by the onset of one of the tactile stimulation patterns. The LEDs were colored red, blue, green, and yellow (shown in **Figure 1F**) in order to cover as wide a range of the wavelength color spectrum as possible because this was expected to increase the number of activated photoreceptors.

### Neuropixel recordings and extraction of neurons

We recorded spiking activity across somatosensory (S1) cortical structures with Neuropixel silicon probes. S1 recordings were made in the forepaw region with coordinates −1.0 – 1.0 mm relative to bregma and 3.0-5.0 mm lateral to the midline, contralateral to the stimulated paw. The Neuropixel probe is 10 mm long and has a total of 384 channels for recording. The probe was inserted all the way and recorded from all channels but only the first 100 channels were used for analysis as these were located in the approximately 1.5 mm deep cortex of the rat.

Neuropixel recordings were processed with the Kilosort2.5 Matlab package for spike sorting. All units detected by the software were visually inspected in the open-source Python library Phy and selected based on their shape, spike frequency, amplitude, and behavior. Units with average amplitudes lower than 1μV and units with spike frequencies lower than 0.5 Hz were automatically deselected. Stimulation artefacts incorrectly classified as neural units by Kilosort were identified by their shape and then manually deselected. Units which had inter-spike-intervals (ISI) around 0 ms (an example ISI histogram is shown for one sample neuron in **Figure 1C**) were regarded as multiple units because of violation of the refractory period limiting the firing activity of any single neuron. If the ISI plots could be improved by splitting the unit into two or more, the units were kept. If the plots did not improve, the units were deselected and not included in the analysis.

### Spike response decoding

To estimate decoding accuracy in individual neurons we analysed evoked spike trains from individual neurons. Spike trains were extracted in the time window between stimulation onset and 600 ms post stimulation onset and convolved into a continuous vector with a gaussian kernel of 5 ms. Convolved spike trains were z-scored before being randomly split into a test and a train data set with equivalent numbers of spike train responses to individual stimulation types. We used principal component analysis (PCA) to extract dimensions explaining 95% of the variance across the z-scored spike responses in the train set. To determine how each of the spike responses were placed in PC space, we calculated the score for each PC relative to each of the z-scores responses in the train set and the test set. A kNN classification algorithm with 9 neighbors was trained on the scores from the train set before classifying scores from the test set as belonging to one the stimulation patterns and was repeated 50 times. The level of uniqueness in the temporal response profile to a stimulation pattern would determine the accuracy with which the algorithm could classify it.

### Decoding analysis 8×8

The 8×8 decoding analysis included responses for all four tactile stimulation patterns with and without simultaneous visual stimulation. This analysis was performed to get an estimate of the extent to which individual neurons responded in distinct ways to the activation patterns depending on whether visual stimulation was also present. As a measure for decoding accuracy in the 8×8 classification analysis, we used the F1 score. First, precision and recall were calculated with True Positives (TP), False Positives (FP) and False Negatives (FN):

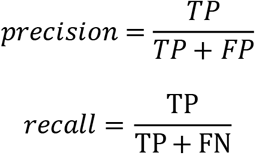

With the precision and recall parameter for each of the 8×8 matrices, the F1-scores were calculated:

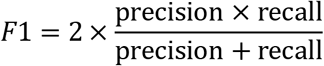

We redid the 8×8 decoding analysis, this time with shuffled labels for all spike responses. Training the algorithm on a data set with shuffled labels result in random guessing and plotting the distribution of all F1 scores for both the shuffled and the non-shuffled 8×8 decoding analyses is a way to ensure that the F1 scores in the non-shuffled classification are not due to chance.

### Separation individual patterns

To estimate how well the algorithm had separated the presence and absence of visual information for the individual patterns, 2×2 matrices were extracted from the 8×8 matrices (see example in **Figure 3A** and **Figure 3C**). Each of the extracted 2×2 matrices were normalized to 1 and the diagonal sum of each matrix was used as a measure of the accuracy with which the algorithm had classified responses to individual patterns with and without visual stimulation. A diagonal sum above 0.5 in the normalized matrices would mean that the algorithm had more correct guesses than incorrect guesses.

**Figure 2.**
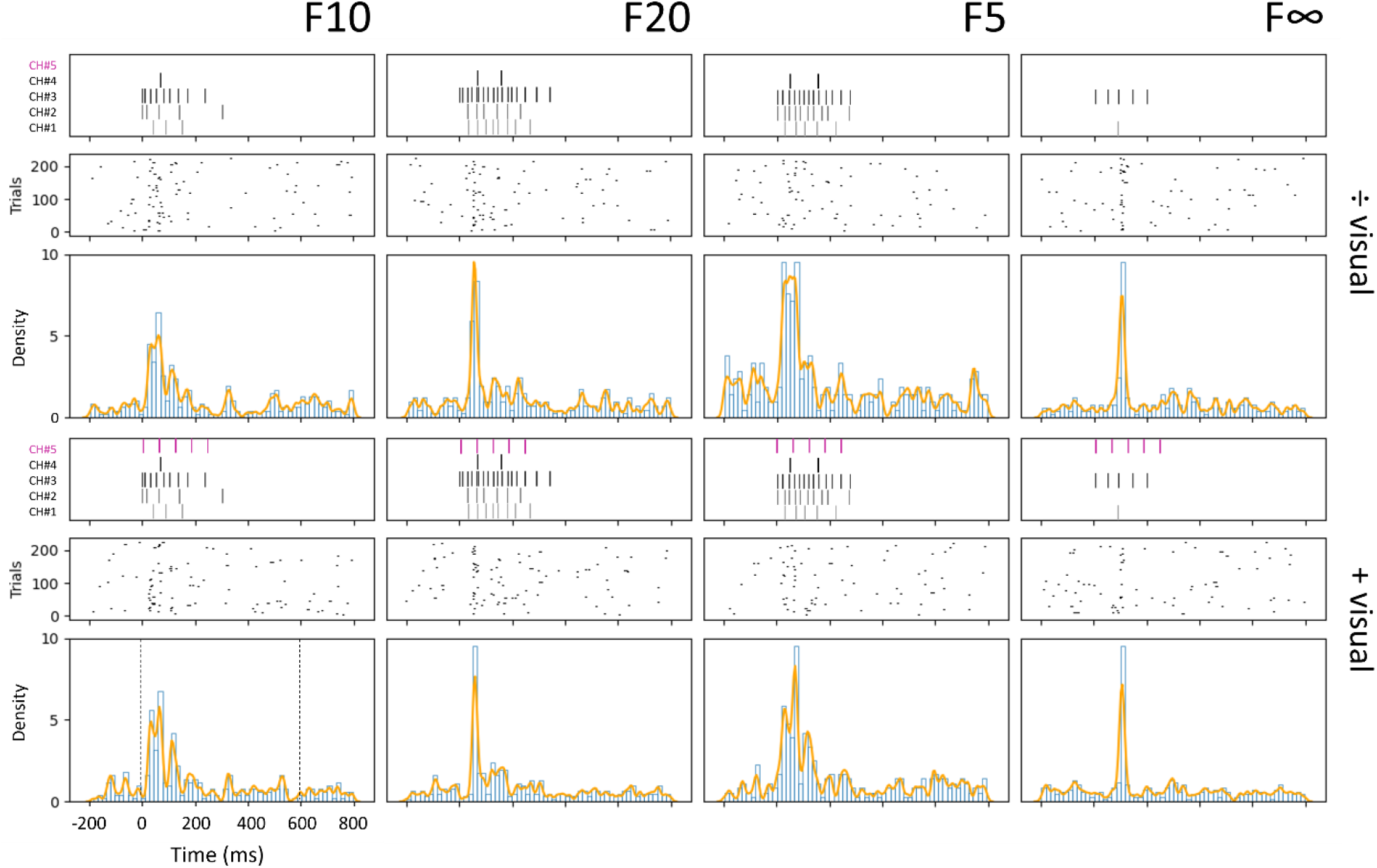
Illustration of the stimulus combinations (the 4 tactile stimulation patterns with or without the visual stimulus) and responses of a sample S1 neuron. From the top: Illustration of the 4 spatiotemporal activation patterns without visual flashes (top row) and with visual flashes (bottom row). Below each respective stimulation pattern are raster plots and the PSTHs (with superimposed KDE curves) of the spike responses from one sample neuron. The dashed lines in the lower left plot shows the time window used for the kNN classification analysis (0-600 ms post stimulus onset).

**Figure 3.**
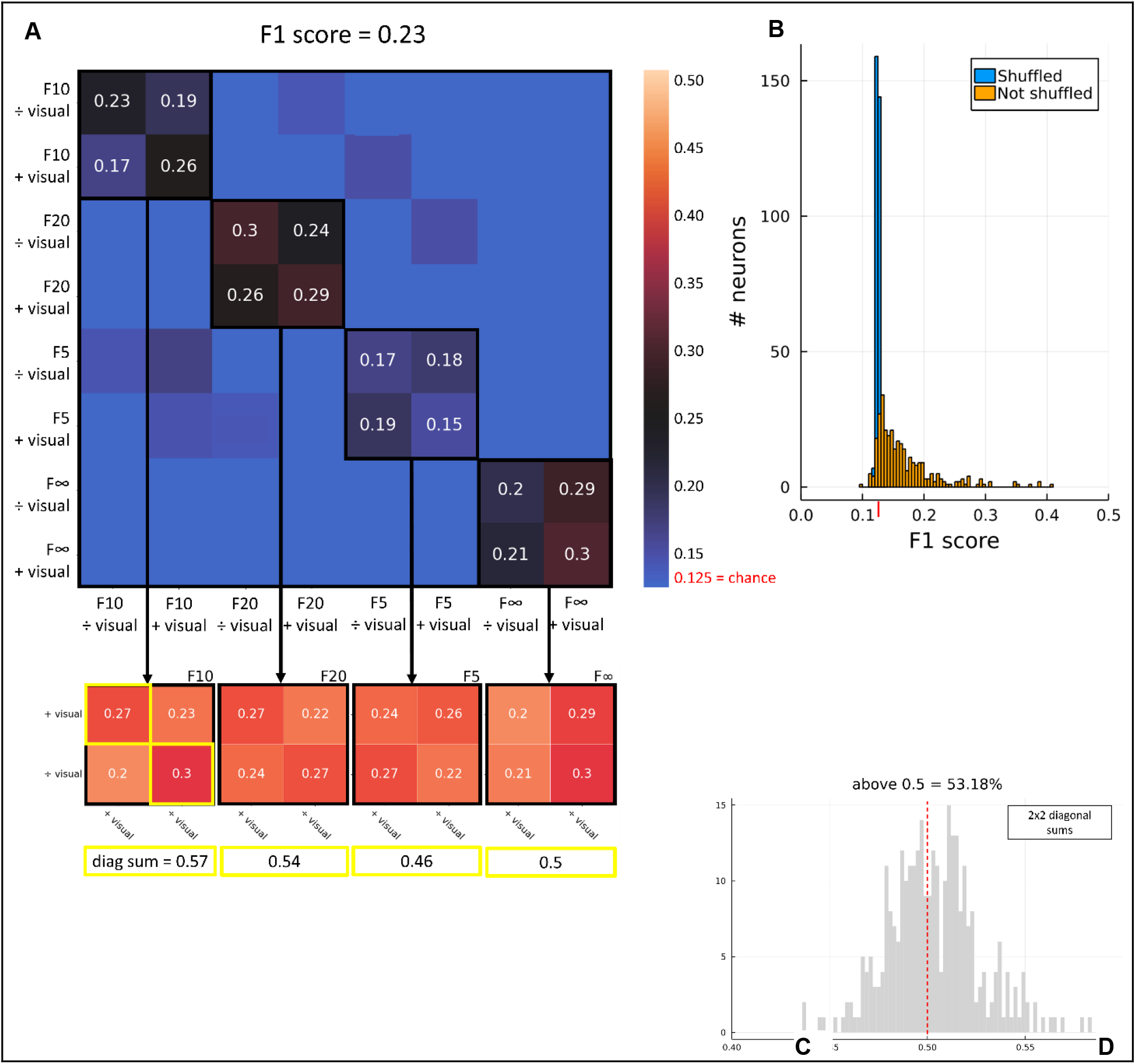
Decoding performance at single neuron level **(A)** 8×8 decoding of all tactile patterns with and without visual stimulation for one sample neuron. **(B)** Distribution of F1 score of the 8×8 decoding analysis, shuffled and not shuffled. **(C)** Normalized 2×2 matrices extracted from the 8×8 matrix above. Below the diagonal sum of each of the matrices. **(D)** Distribution of 2×2 diagonal sums from all 8×8 decoding matrices.

### Tactile and visual decoding

A non-responding neuron or a neuron with non-distinctive responses to the tactile stimulation patterns will not have uniquely modified responses to visual stimulation. Therefore, we wanted to analyze if there was some relationship between decoding accuracy of tactile patterns and decoding accuracy of patterns with and without visual stimulation. We performed a 4×4 kNN classification analysis to get an estimate of the neurons’ level of tactile decoding. In this analysis, the algorithm only had to classify spike responses to the four tactile activation patterns without visual stimulation. As in the 8×8 decoding analysis, we did the 4×4 decoding with and without shuffled labels. From the shuffled 4×4 decoding we calculated the mean F1 score and the standard deviation (SD). We next extracted six populations of neurons with F1 scores 2, 4, 6, 8, 10, and 12 SD above the shuffled mean respectively. The shuffled mean is theoretically equivalent to chance (0.25). Thus, neurons in these population all had F1 scores above chance and they were considered as positively decoding the tactile patterns with increasing accuracies. For each of these six groups of neurons, we plotted the diagonal sums from the normalized 2×2 matrices to see if the distributions would change as the populations included neurons with increasingly accurate tactile decoding.

### Cooperative decoding analyses

To find the most accurate representation of all stimulation patterns with and without visual stimulation among all spike responses, we performed a cooperative decoding analysis. The idea is that even though the information about the tactile stimulation patterns and the presence or absence of visual stimulation is expressed at the level of individual neurons, it is more accurately expressed in a population of neurons. The cooperative decoding analysis for the 8×8 classification was performed by training the kNN classifier with spike responses for ascending groups of neurons by adding one neuron at a time, starting with the best performing neurons ordered by their F1 score in the individual 8×8 matrices. The largest analyzed population was set to 16 neurons because decoding of larger population was too computationally heavy. For each 8×8 population decoding analysis, the 2×2 matrices were extracted and normalized, and the diagonal sums were used as a measure of improvement for decoding of visual stimulation for individual patterns.

## Results

Our aim was to investigate the effect of visual stimulation on S1 neuronal responses to different types of naturalistic tactile stimulation. For this purpose, we used four different spatiotemporal tactile stimulation patters delivered with and without simultaneously flashing multicolor LEDs. The tactile stimulation patterns mimic the activation of primary afferents at local skin sites when touching surfaces with different types of roundness. We recorded the spike activity from neurons in the forepaw region of primary somatosensory cortex (S1) with a Neuropixel probe in 4 anesthetized rats. Neurons were automatically identified with the Matlab package Kilosort2.5 and visually inspected in the open-source Python library Phy. A total number of 453 neurons were selected for plotting. After plotting, 137 units were discarded because they were mixed with artefacts, had too low spike activity or had too much noise. A final number of 317 units were included in the analysis of the spike responses evoked by the 4 tactile stimulation patterns with and without simultaneous visual stimulation.

### Decoding of inputs for individual neurons

The spike responses to the four spatiotemporal activation patterns with and without simultaneously delivered visual flashes was first explored by visually inspecting raster plots and peristimulus time histograms (PSTH) (**Figure 2**). The PSTHs were overlaid with a Kernel Density Estimation (KDE), which represents each spike event as a Gaussian distribution to minimize the impact of the discretizing step present in the PSTHs due to binning. The KDE thus transforms the temporal spike responses from discrete events to a continuous function. Each of the KDE curves overlaid the eight PSTHs (four tactile patterns +/− visual stimulation) correspond to the average of the convolved spike responses for that stimulus condition (see Materials and Methods for a more detailed description of convolution). **Figure 2** displays the raster plots and PSTHs for one sample neuron. Although there appears to be a lot of variation in the spike responses across the 200 repetitions of each of the tactile patterns +/÷ visual stimulation, some clustering is visible around the onset of each stimulation as we have also seen in previous raster plots for the input patterns (Enander et al., 2019; Oddo et al., 2017). The variation is less visible in the PSTH where the KDE curves show substantial differences between the four tactile patterns and smaller differences between the individual patterns depending on the presence of visual stimulation.

To quantify the uniqueness of the responses to the patterns with and without visual stimulation we first completed an 8×8 classification analysis for each individual neuron (**Figure 3**). The classification analysis was based on plotting the position of each single convolved spike response (one for every repetition of the stimulus presentation) in a principal component space. Then, the classification algorithm detected if the individual response was located closest to the responses of the same stimulation type, or those of other stimulation types. The kNN classifier in this way obtained a measure of the accuracy of the classification across all the responses evoked by the same type of stimulation (see Materials and Methods for a more detailed description). **Figure 3A** illustrates the decoding performance for one sample neuron. Notably, there are higher classification scores in the diagonal, which indicates that the responses evoked by the same tactile input patterns were more similar to each other than to the responses evoked by other tactile input patterns. Higher values in the 2×2 blocks along this diagonal indicate that also the responses evoked by the same tactile input patterns, regardless of whether they were delivered with or without simultaneous visual input matrix, were also more similar to each other. Finally, the higher values in the diagonal within these 2×2 blocks indicate that this cell had somewhat different responses whether the given tactile input pattern was delivered with or without simultaneous visual input.

The confusion matrix in **Figure 3A** reveals higher diagonal values in the highlighted 2×2 squares for the patterns F10 and F20 compared to the non-diagonal values. This is an indication of accurate classification of both F10 +/÷ visual and F20 +/÷ visual. In the highlighted 2×2 squares for F5 +/÷ and F∞ +/÷ visual, the diagonal values are not higher than in the opposite diagonal, indicating that the neuron responses in these two cases did not provide information by which the condition +/÷ visual could be separated. This may be due to the differences not being consistent enough for the classifier to accurately categorize the spike responses. We have previously seen these “preferences” for neurons to respond more distinctively and consistently to some tactile patterns, resulting in high decoding accuracy for some patterns and low decoding accuracy for other patterns (Enander et al., 2019; Oddo et al., 2017). We will discuss this in more detail in the Discussion section.

The 8×8 matrix in **Figure 3A** also shows the F1 score for the sample neuron across all conditions; 0.23. This is clearly above the theoretical chance level, which was 0.125. The distribution of all F1 scores in the 8×8 decoding analysis can be seen in **Figure 3B** (orange). We also shuffled the stimulus labels across the responses to see what the chance level would be given that the neuron spike responses could have some intrinsic structure which made it easier for the classifier to make spurious classifications. The distribution of F1 scores in the shuffled 8×8 decoding analysis is shown in the same figure (blue). It can be seen that the chance level obtained after response shuffling was tightly located around the theoretical chance level of 0.125. Across the whole population of S1 neurons included in the 8×8 classification analysis, a total of 286 neurons had an F1 score above chance (0.125).

Notice that although the F1 score in the 8×8 matrix in **Figure 3A** is above chance, only two of the patters in the 2×2 matrices have a diagonal sum above 0.5. Because the F1 score does not give insight into the accuracy with which the algorithm separated responses to individual patterns with and without visual stimulation, we next extracted all of the 4 2×2 matrices obtained from each 8×8 matrix (one per neuron). **Figure 3C** displays normalized 2×2 matrices extracted from the 8×8 matrix in **Figure 3A**. Written below each of the 2×2 matrices is the sum of the diagonal. We next started to use this measure, to quantify the differences between responses with and without visual stimulation. **Figure 3D** shows the distribution of all 2×2 diagonal sums from all 8×8 decoding matrices. The red dashed line indicates the chance level of 0.5. As can be seen from this plot, the average separation level was >0.53, indicating that the responses obtained by the different tactile input patterns could with a probability somewhat higher than chance be separated based on whether the tactile input was delivered with or without the visual input.

### Relationship between tactile decoding and separation of patterns

Since we knew that some neurons did not separate the tactile input patterns well to begin with (**Figure 2B**), we next sorted the neurons based on how well they could separate the tactile input patterns. Neurons without a clear response to the tactile input could not be expected to separate well the condition, i.e. whether the tactile input was delivered with or without the visual stimulation. Therefore, we first analyzed the F1 score for each neuron looking only at the four tactile input patterns without visual stimulation (**Figure 4A**). From this plot, we also obtained a standard deviation of the shuffled responses (blue bars in **Figure 4A**), i.e. the standard deviations of the actual chance level given the structure of the spike responses themselves. We could then repeat the 2×2 diagonal sum calculation from above (**Figure 3C**) but now only including neurons with a certain performance for the tactile input patterns alone, quantified as the numbers of standard deviations away from the chance level (**Figure 4B**). The six populations are neurons with an F1 score in the tactile decoding analysis of 2, 4, 6, 8, 10, and 12 SD above chance (0.25). (green lines in **Figure 4A**). The plot to the far left in the top row of **Figure 4C** shows the distribution of the neurons with an F1 score 2 SD above chance. The threshold is increasing in the plots towards right with the highest threshold of 12 SD above chance in the right bottom row. 259 neurons had an F1 score 2SD above chance, 240 4 SD above chance, 218 6 SD above chance, 203 8 SD above chance, 180 10 SD above chance, and 164 12 SD above chance. When plotting the distribution of the 2×2 diagonal sums for neurons in the different populations, it can be seen that an increasing number of the 2×2 diagonal sums fall above chance of 0.5, starting with 53.67% for the population of 259 neurons and finishing with 57.32% for the population of 164 neurons. This shows that of all recorded neurons, neurons with higher decoding performance in the 4×4 analysis also had higher accuracy scores in the 2×2 analysis, indicating that at least to some extent neurons with higher tactile decoding also are more likely to respond differently when visual stimulation is also present.

**Figure 4.**
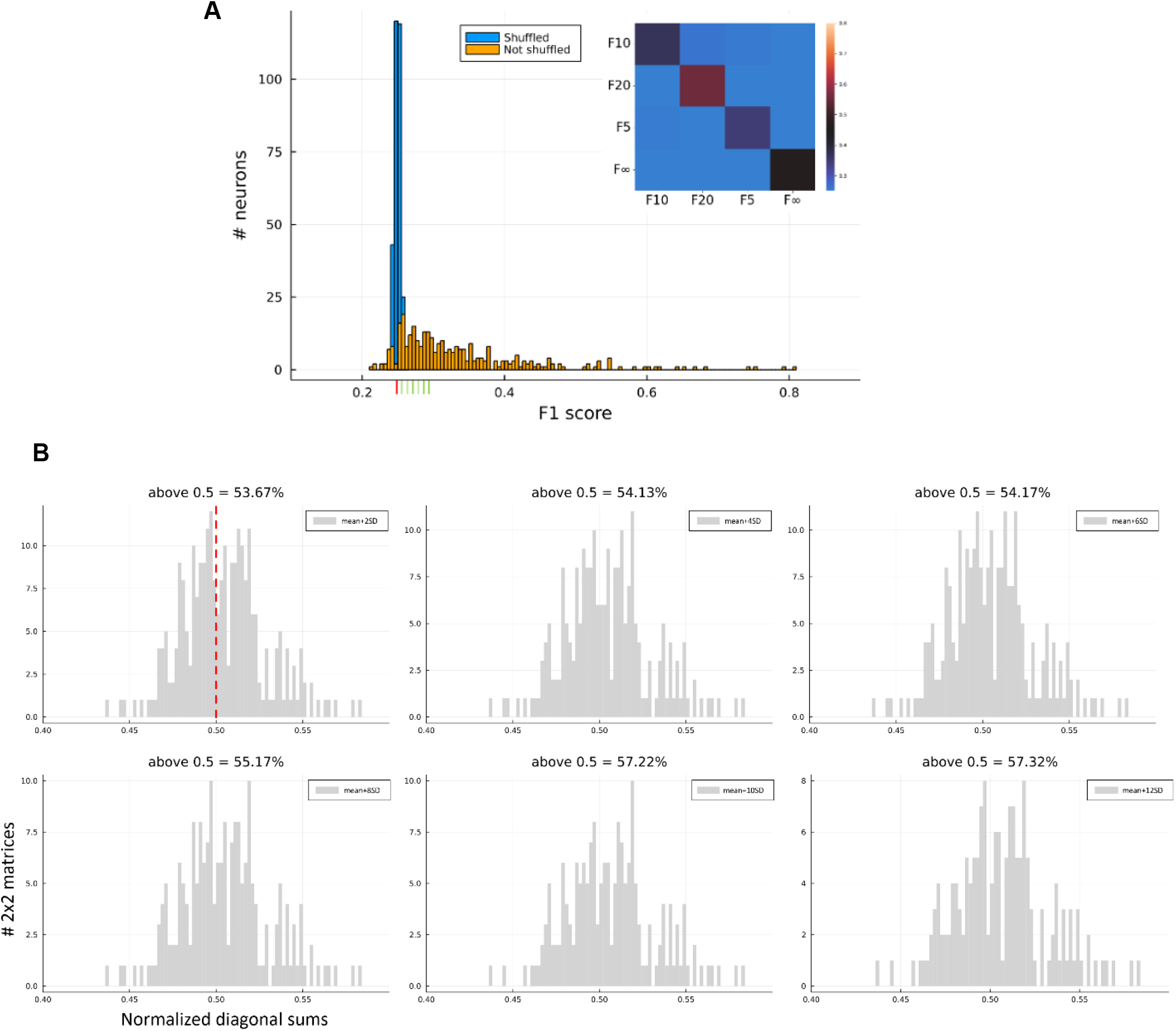
Decoding performance at single neuron level **(A)** Distribution of F1 scores in the 4×4 decoding analysis of the four tactile stimulation patterns, shuffled (blue) and not shuffled (orange). The red line in the bottom indicates the mean F1 score in the shuffled analysis and the 3 green lines indicate the mean + 2, 4, 6*, 8, 10, and 12* SD. In the right corner of the figure is the 4×4 classification matrix for the same sample neuron as in *F*igure 3A. **(*B*)** Distribution normalized 2×2 diagonal sums for *six populations of* neurons*. Populations from top left towards right: Neurons with* a *4×4 decoding* F1 score *above the mean + 2,* 4, 6, 8, 10, and 12 SD.

### Complementary decoding of inputs

It has previously been shown that combining spike responses from a population of neurons enhances classification accuracy of tactile input patterns (Enander et al., 2019; Oddo et al., 2017). In these studies, individual S1 neurons allowed for co-operative decoding, indicating that each neuron represented specific aspects of the tactile input. We wanted to test if combined responses from a population of S1 neurons similarly would improve decoding of tactile input patterns with and without visual flashes.

After creating a version of the classification analysis that could take into account spike responses from populations of neurons, we ran the 8×8 classification analysis for a growing population of neurons. We started with spike responses from a population consisting of the two neurons with the highest F1 scores from the individual 8×8 decoding analysis. We then ran the 8×8 decoding for the population, added the next neuron listed by its F1 score, ran the decoding analysis and so forth. We continued the process until we reached a population of 16 neurons (**Figure 5A**). Each time we ran the population classification analysis we extracted and normalized the 2×2 matrices in the same way as in the individual classification analysis. The sums of the 2×2 diagonals for the growing population of neurons are displayed in **Figure 5B.** Here it can be seen that for the population of two neurons, separation of the patterns F10, F20 and F5 +/÷ visual was above chance (0.5) whereas separation of F∞ +/÷ visual was below chance. The decoding of F∞ +/÷ visual reaches above chance decoding for the population of 7 neurons.

**Figure 5.**
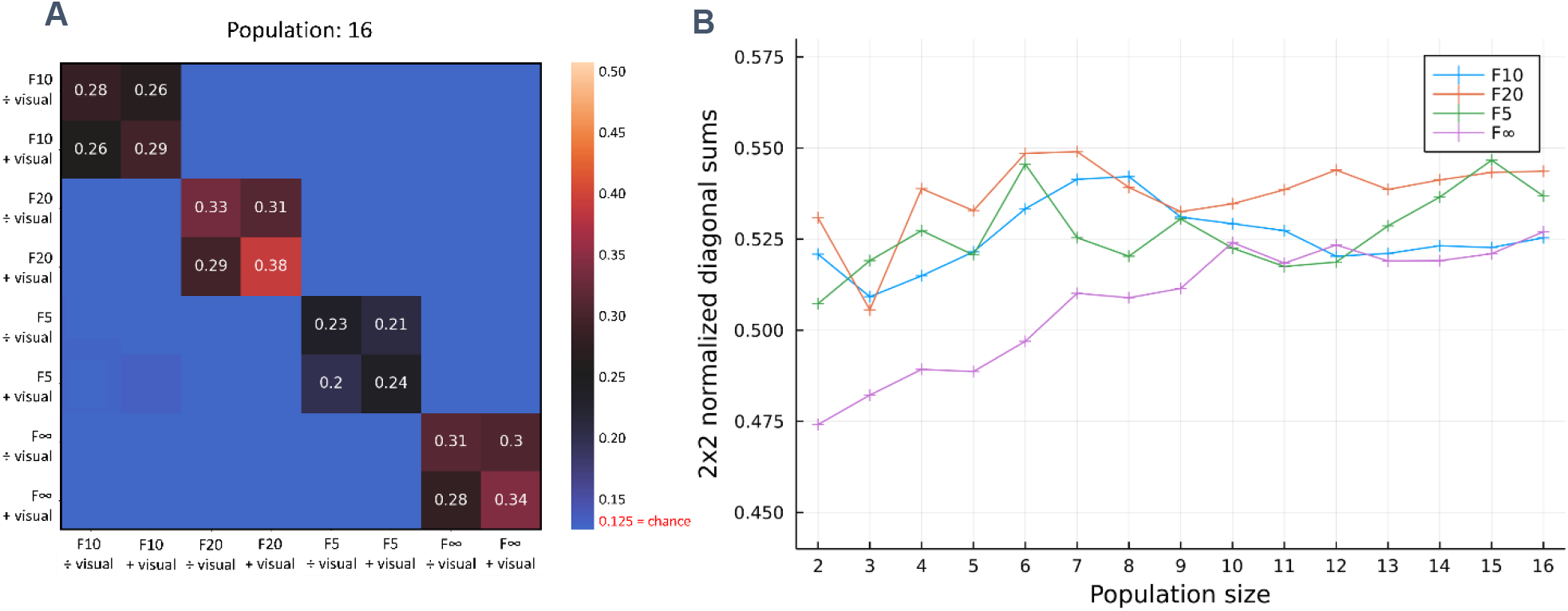
Decoding performance at population level. **(A)** 8×8 heatmap for population decoding of 16 neurons. **(B)** Diagonal sums of 2×2 matrices extracted from 8×8 classification analyses for populations 1-16 neurons.

## Discussion

We found that spike responses to tactile activation patterns delivered to the distal digit 2 of the rat forepaw are weakly but consistently modulated by simultaneous visual flashes in S1 neurons. Our study thus supports previous observations that perturbation of the activity in remote cortical areas, i.e. focal stroke-like lesions (Wahlbom et al., 2019) or local circuitry perturbation through intracortical electrical microstimulation (Etemadi et al., 2022) have profound effects on the information processing of tactile input patterns in neurons of the forepaw region of somatosensory cortex. While these observations are all compatible with that for example visual input would have profound impact on the cortical circuitry representations of given tactile input patterns (and vice versa), i.e. visual input would impact the perception of those given tactile input patterns, it could be argued that the manipulations used so far to demonstrate the cortical circuitry behavior were artificial and not reflective of normal information processing in the neocortex. Here we show that natural external stimulation also impacts the interpretation of given tactile input patterns indicating that the cortical network may be far more dynamic in its processing than traditionally thought. As previously suggested (Etemadi et al., 2022) the specific “network solution”, i.e. neuronal response to a given input, depends on the interplay between the global state of cortex and the actual sensory input. We believe that flashing lights impact the global cortical state and affect the network solutions to tactile input which in turn result in the output in S1 neurons being modified.

### Potential limitations of the setup

We wanted to investigate the dynamic modulation of spike responses and avoid unnecessary spontaneous activity that arises in awake animals. Any spontaneous activity will in the analyses be equal to noise and subtle modulations of responses would be harder to detect. Along with this objective comes the challenge of having visual stimulation that in any way can be meaningfully matched with tactile stimulation, the problem being that the semantic properties of congruent visuo-tactile stimulation will not be perceived by an animal that’s asleep. A second challenge is that the focal plane is drifting under anesthesia causing the focus of the eye to change. This would have implications if video or images were used with the aim of having more naturalistic visual stimulation. In any case, we believe that we have shown that multimodal properties of neurons extend beyond integration of congruency of multiple sensory input.

Similar to previous studies using electrotactile spatiotemporal input patterns (Enander et al., 2019; Etemadi et al 2022; Norrlid et al., 2021; Oddo et al., 2017; Wahlbom et al., 2021) the approach was motivated by having both reproducible and naturalistic skin sensor activation. The variability that occurs through natural mechanical activation of mechanoreceptors inevitably also causes variation in the cortical representation of tactile input, making it harder to experimentally detect similarities in spike responses. With spatiotemporal electrical stimulation we overcome this variability while still having stimulation that evolve in space and time as natural mechanical stimulation would do (see Oddo et al., 2017 and Enander et al., 2019 for a more detailed discussion of the tactile input patterns). Although the electrical tactile stimulation was mild, it is still a direct stimulation of the peripheral nerve fibers, and it is possible that it is inducing stronger synaptic input to the cortex than the visual stimulation. Therefore, it is not unlikely that responses to natural mechanical stimulation in S1 neurons would be even more modulated by flashing lights.

### What we know about multimodal processing

Although many neurophysiology studies have revealed responses to a variety of modalities in primary sensory regions of the cortex (Alsius et al., 2005; Busse et al., 2005; Enander et al., 2019; Koelewijn et al., 2010; Talsma & Woldorff, 2005; Wallace et al., 2004; Zhou et al., 2020), very few studies have investigated the principles by which integration or modulation of multiple sensory input occur at the single neuron level. Of those that did, some focussed on the semantic aspects of multisensory integration by recording from neurons in macaque monkeys engaging in behavioral tasks (Alsius et al., 2005; Barraclough et al., 2005.; Zhou et al., 2020). The common approach in these studies is a comparison of mean spike density frequencies to unimodal vs multimodal input, for example videos of different actions shown with and without sound. While there undeniably is value in having this type of natural stimulation in the experiments, unfortunately the rich information in the temporal evolution of the spike responses to different types of time-evolving input is lost by reducing temporal response profiles to means. We believe that cortical processing of multimodal sensory information is dynamic and should be studied as such.

## Acknowledgements

This work was supported by the EU H2020 MSCA Grant project# 861166, ‘INTUITIVE’

